# Association Analysis of *PAEP*, *KRT*10 and *BMP*7 Genes SNPs with Reproductive Traits in Kele Pigs

**DOI:** 10.1101/2024.09.14.613052

**Authors:** Yong Zhao, Chunyuan Wang, Yan Wu, Jin Xiang, Yiyu Zhang

## Abstract

The aims of this study were to investigate the effects of single nucleotide polymorphisms (SNPs) in progestogen-associated endometrial protein (*PAEP*), keratin 10 (*KRT*10), and bone morphogenetic protein 7 (*BMP*7) genes on reproductive traits (total number of piglets born, number of piglets born alive, litter birth weight, number of piglets weaned and litter weight weaned) in Kele pigs. We used a total of 255 multiparous Kele sow (2-4 parities) as research materials. SNPs were identified by PCR amplification instrument and sequence alignment software DANMAN. The population genetic characteristics of SNPs were analyzed using the SHEsis online software. Bioinformatics analysis of SNPs were conducted using RNAfold, SOPMA, SWISS-MODEL, and Swiss-PdbViewer programs. The associations between the SNPs and reproductive traits in Kele pigs were analyzed through SPSS 22.0 software. In this study, nine SNPs were identified in the three genes: g.1884992 T>C (exon 4), g.1885152 G>C (intron 4) and g.1887834 G>A (intron 4) in *PAEP*, g.21643703 C>T (intron 4), g.21643714 G>A (intron 4) and g.21643741 G>A (exon 5) in *KRT*10, and g.57647887 G>A (intron 3), g.57647990 C>T (intron 3) and g.57648145 C>G (intron 3) in *BMP*7. In SNPs g.1884992 T>C of *PAEP,* missense mutation eventuated structural changes in mRNA and proteins secondary structure. In SNPs g.21643741 G>A of *KRT*10, synonymous mutation led to alteration in mRNA secondary structure. For *PAEP,* the CC genotype in SNPs g.1885152 G>C and the AA genotype in SNPs g.1887834 G>A showed significantly higher values than other genotypes in all reproduction traits except for litter birth weight, preliminarily identified as favorable genotypes. For *KRT*10, the GG genotype in SNPs g.21648641 G>A showed significant superiority than AA genotype (*P<*0.05) in all aspects except for litter birth weight, and notably surpassed the GA genotype in total number of piglets born (*P<*0.05), preliminarily recognized as a favorable genotype. Regarding *BMP*7, the GA genotype in SNPs g.57647887 G>A and the CT genotype in SNPs g.57647990 C>T exhibited significantly higher number of piglets born alive and number of piglets born alive compared to other genotypes (*P<*0.05). And the GG genotype in SNPs g.57648145 C>G was significantly associated with higher litter birth weight (*P<*0.05). The result of diplotype analyses indicated that the H3H3 (CCGGGG) of *PAEP* and H3H3 (CCGGAA) of *KRT*10 had a significant effect on the five traits. For *BMP*7, the H4H4 (AATTGG) diplotype showed a significant influence on all genotypes except litter birth weight.

## Introduction

Kele pigs, native to the Yunnan-Guizhou Plateau, have excellent meat quality with strong water retention, juicy muscle, good color, and is an excellent ingredient for Xuanwei ham. However, they are accompanied by poor reproductive traits, such as low litter size and teat number [1] [2]. Reproductive efficiency has a great impact on the economic benefit of pork production [3]. As the improvement of total living standards, there is an increasing global demand for pork, which precipitates the pig farming industry has been more and more prosperous. The poor reproductive traits of Kele pigs has been hard to meet the demands of pork yields in modern society, which severely impact their development. Therefore, improving the reproductive traits of Kele pigs holds practical significance for enhancing the efficiency and sustainability, increasing production efficiency, and meeting market demands.

*PAEP*, also known as β-lactoglobulin (*BLG*) gene, spans 4193 bp including 7 exons and 6 introns. The protein encoined by this gene consists of 180 amino acids and belongs to the Nuclear Lipid Calcium-Binding Proteins Family [4]. As a secretory protein usually expressed in the reproductive tract with anti-tumor growth characteristics, PAEP has been primarily studied in reproductive system diseases [5][7]. In addition, β-lactoglobulin coding genes often have a favourable effect on milk traits. A large number of authors have reported the study of *BLG* polymorphism on milk production traits and milk quality traits [8] [9], only a few research reports have been published regarding their association with reproductive traits. Felenczak et al. [10] found there was a tendency towards lower age at first calving and longer calving interval in Simmental cows with the β-lactoglobulin gene BB genotype. Tsiaras et al. [11] found Holstein cows with AA genotype had significantly shorter gestation length than did those with AB or BB genotype. However, its polymorphism has not been reported in the reproductive traits of pigs.

*KRT*10, located on chromosome 12 of pig, with 8 exons and 7 introns, is a gene encoding keratin 10 (K10). Keratin 10 is a type I keratin, as the main structural protein of the epidermis [12]. Nowdays, the research on KRT10 has been focused on various disease mechanisms [13] [14], especially the regulation of ichthyosis [15]-[18]. In addition, scientists try restoring the mutaion of *KRT*10 gene to restore normal skin structure and function by gene therapy or cell engineering techniques [19] [20]. The studies have been proved that *KRT*10 also has potential expression regulation on the reproductive physiology of livestock and poultry. Smolenski et al. [21] found that several members of the keratin family, including *KRT*1, *KRT*4, *KRT14*, *KRT*17, and *KRT*10, participate in the immune defense against pathogens in infected bovine mammary glands. In the study by Gao et al. [22], the downregulation of *KRT*10 gene expression may be related to the development of sheep mastitis. Cunha et al. [23] found KRT10 appears after endogenous estrogen converts the vaginal epithelium into glycogen squamous epithelium, and is a marker of terminal differentiation of squamous epithelium. In view of this, although *KRT10* has not been studied in terms of polymorphism, it is worth pioneering research on the association between its polymorphism and reproductive traits.

*BMP*7, also known as bone morphogenetic protein-7, is a member of the transforming growth factor beta (TGF-β) supergene family of functional proteins [24], and it was identified by Celeste et al. [25]. *BMP*7 can induce undifferentiated mesenchymal cells in the perivascular and connective tissues to differentiate into bone and cartilage cells, leading to the formation of bone tissue, playing a crucial role in the regulation of bone homeostasis and vertebral development [26] 2[27]. *BMP*7 is expressed in ovary, uterus and mammary gland of livestock and poultry. Scientists have suggested that *BMP*7 may be essential for normal folliculogenesis and ovulation [28] [29]. Furthermore, The decrease in *BMP*7 expression may facilitate decidulization of the endometrium, thus aiding the establishment of pregnancy (**^Error! Reference source not found.^**. In the study of **Error! Reference source not found.** suggested that *BMP*7 may antagonize transforming growth factor β 1 (TGFβ1) in tumorigenesis-associated epithelial-mesenchymal transition (EMT) in breast cancer, so BMP7 may be a promising therapeutic target for treating breast cancer. *BMP*7 also exhibits high expression levels in cervical cancer, making it a potential therapeutic target for cervical cancer treatment (^Error! Reference source not found. 加多态的东西^

Based on regulatory significance of *PAEP*, *KRT*10 and *BMP*7 in reproductive physiology, we tried to analyzed the associations between SNPs and reproductive traits in Kele pigs, aiming to provide genetic markers for the reproductive traits of Kele pigs and expedite breeding progress.

## Materials and Methods

### Experimental materials

The experimental pig population was selected from the Kele pig farm in Hezhang County, Guizhou Province, China. A total of 255 healthy multiparous sows (2-4 parities) were randomly chosen, and approximately 0.5 g of ear tissue was collected from each sow for extracting DNA. We also recorded the reproductive statistics of these 255 pigs: total number of piglets born, number of piglets born alive, litter birth weight, number of piglets weaned and litter weight weaned of the first birth. The weaning age of these experimental Kele pigs was 28 days

### DNA Extraction

The DNA was extracted using the Genomic DNA Extraction Kit (Dalian Bao Biotechnology Co., Ltd, Guiyang, China). The extracted genes were evaluated for purity through 1.5% agarose gel electrophoresis and a Nanodrop 2000c spectrophotometer (Thermo Fisher Scientific Co., Ltd, Shanghai, China). The qualified DNA samples were typically between 1.7 and 1.9. Then the qualified DNA samples were diluted to 150ng/ μL and stored at −20°C for later use.

### Primer Design

The DNA reference sequence of the *PAEP*, *KRT*10 and *BMP*7 genes were obtained from the NCBI database (accession number: NW_018084833.1, NC_010454.4, NC_010459.5). Primers were designed using the online software Premier 5.0 (Table 1). All designed primer sequences were synthesized by Shenggong Biotechnology Co., Ltd (Shanghai, China), then stored at −20°C for later use.

**Table 1.**
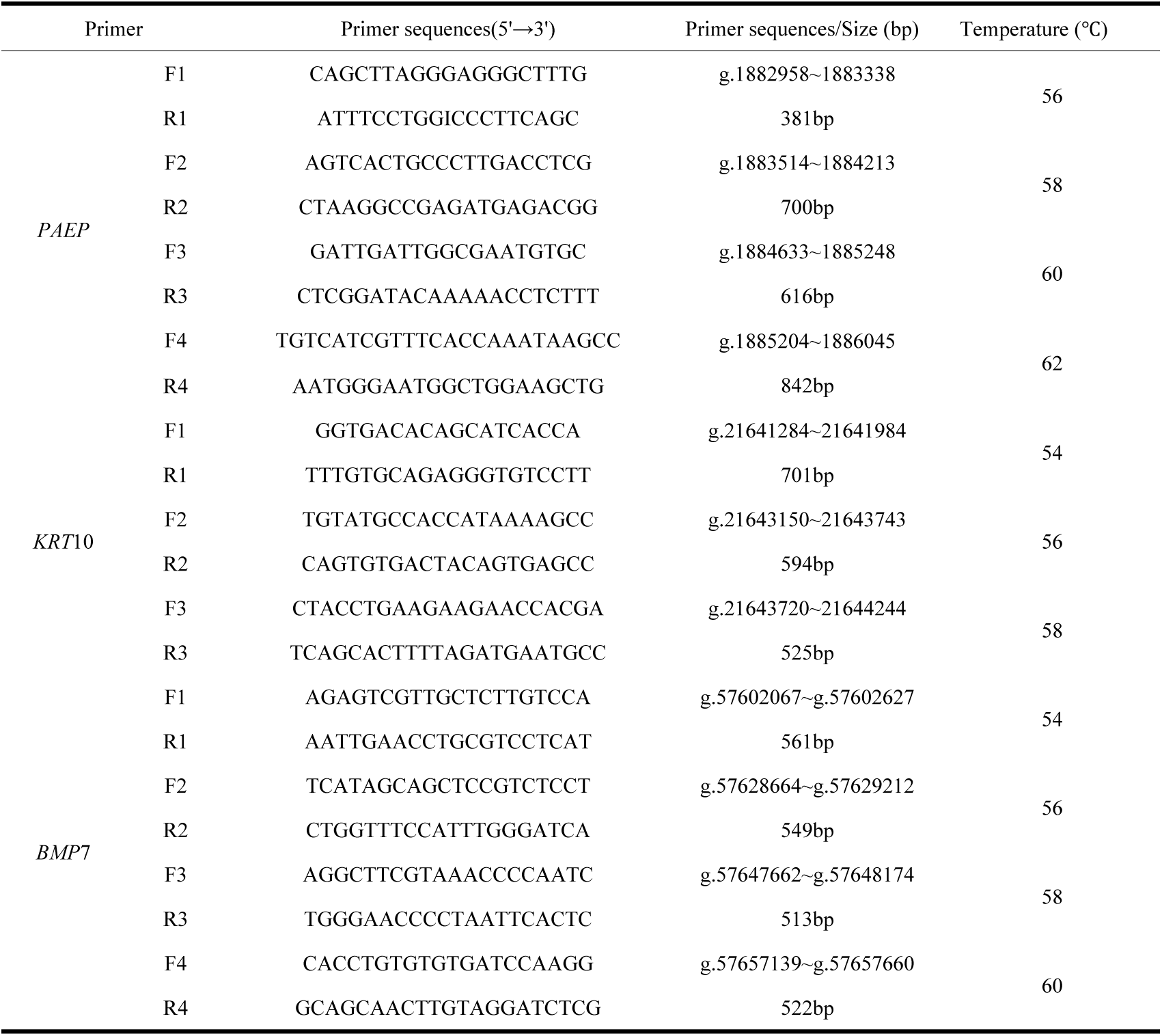
primer sequence information of *PAEP*, *KRT*10, *BMP*7.

### PCR Amplification and SNPs Detection

PCR was performed in a total 20 volume of μL including the following: 1 μL DNA, 10 μL 2× Taq PCR Master Mix, 1 μL forward and reverse primers each (10 mol/L), and 7 μL ddH_2_O. The PCR reaction procedure included initial denaturation at 94°C for 8 min, followed by 32 cycles of denaturation at 94°C for 30 s, and annealing at the optimal temperature for 30 s, then extension at 72°C for 30 s, final extension at 72°C for 8 minutes. The PCR products were analyzed via 1.5% agarose gel electrophoresis. PCR products meeting qualified specificity that the gel bands were clear, bright and free of redundant bands were sent to Shenggong Biotechnology Co., Ltd (Shanghai, China) for sequencing. Sequencing results were used for SNPs identification and genotyping througth Chromas, Editseq software, and manual calculations.

### Statistical analyses

Allelic frequency, genotype frequency, heterozygosity (He), polymorphism information content (PIC), number of effective alleles (Ne) and Hardy-Weinberg equilibrium were calculated by EXCEL2010 software. SHEsis online software (http://analysis.bio-x.cn/) was applied to calculating and analyzing linkage disequilibrium (LD) and haplotypes at SNPs. The multifactorial variable analysis of general linear model (GLM) in SPSS 23.0 software was used for the correlations between SNPs and reproductive traits, polyplotypes and reproductive traits. Multiple comparisons were conducted using Duncan’s method. Data results were presented as mean ± standard deviation. Statistical analyses were performed using Student t test. Mutations’ effects on mRNA secondary structure, protein secondary structure, protein tertiary structure, and local structure were analyzed using online tools, including RNAfold (http://rna.tbi.univie.ac.at/cgi-bin/RNA WebSuite/RNA fold. cgi), SOPMA (https://npsa-prabi.ibcp.fr/cgi-bin/npsa_automat.pl?page=npsa%20_sopma.html), and SWISS-MODEL (https://swissmodel.expasy.org/interactive), as well as the Swiss-PdbViewer software.

## Results

### Summary of Mutation Sites

We identified 9 SNPs totally and obtained the following peak maps according to the sequencing results, and the maps were displayed in Fig. 1 - Fig. 3. The SNPs g.1884992 T>C located in the fourth exon, g.1885152 G>C and g.1887834 G>A were detected in the fourth intron of *PAEP*. The SNPs g.21643703 C>T and g.21643714G>C located in the fourth intron, and g.21643741 G>A located in the fifth exon of *KRT*10. The SNPs g.57647887 G>A, g.57647990 C>T and g.57648145 C>G all located in the third intron of *BMP*7. The g.1884992 T>C missense mutation of *PAEP* led a codon to change from GTC to GCC, causing the amino acid encoded at position 135 to transition from Valine (V) to Alanine (A). The g.21643741 G>A mutation in *KRT*10 changed the codon from AAG to AAA, encoding Lysine (K) in both cases, representing a synonymous mutation.

**Fig. 1.**
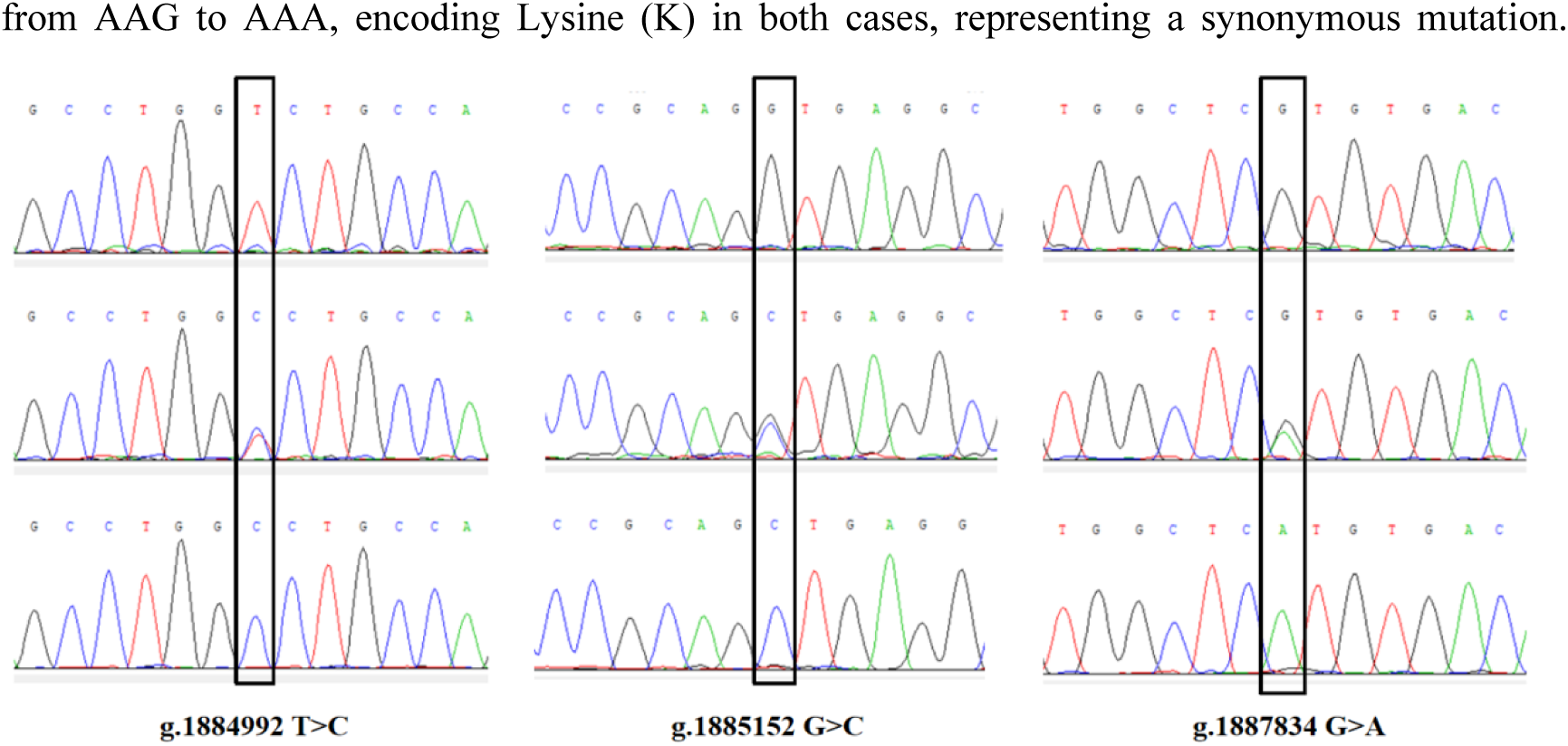
The sequence peak map of the SNPs in *PAEP* gene.

**Fig. 2.**
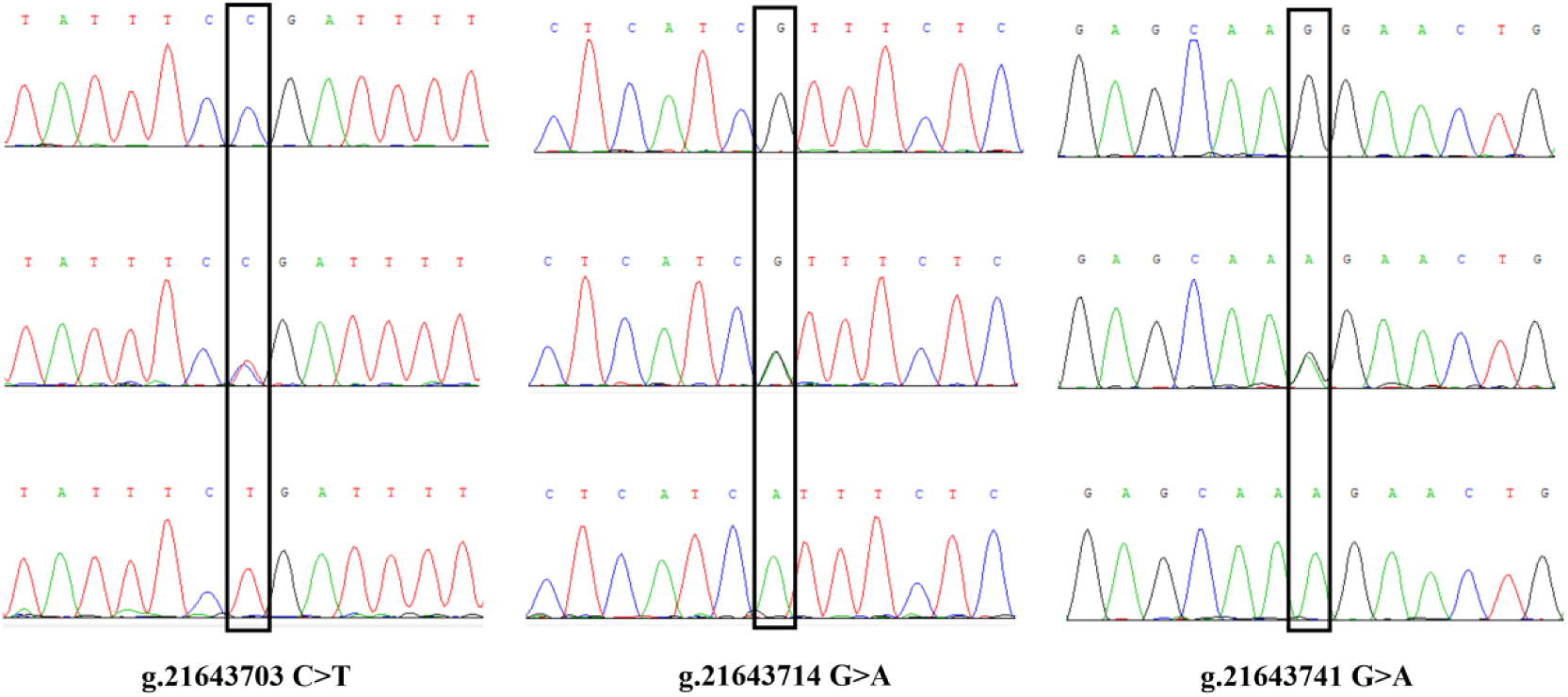
The sequence peak map of the SNPs in *KRT*10 gene.

**Fig. 3.**
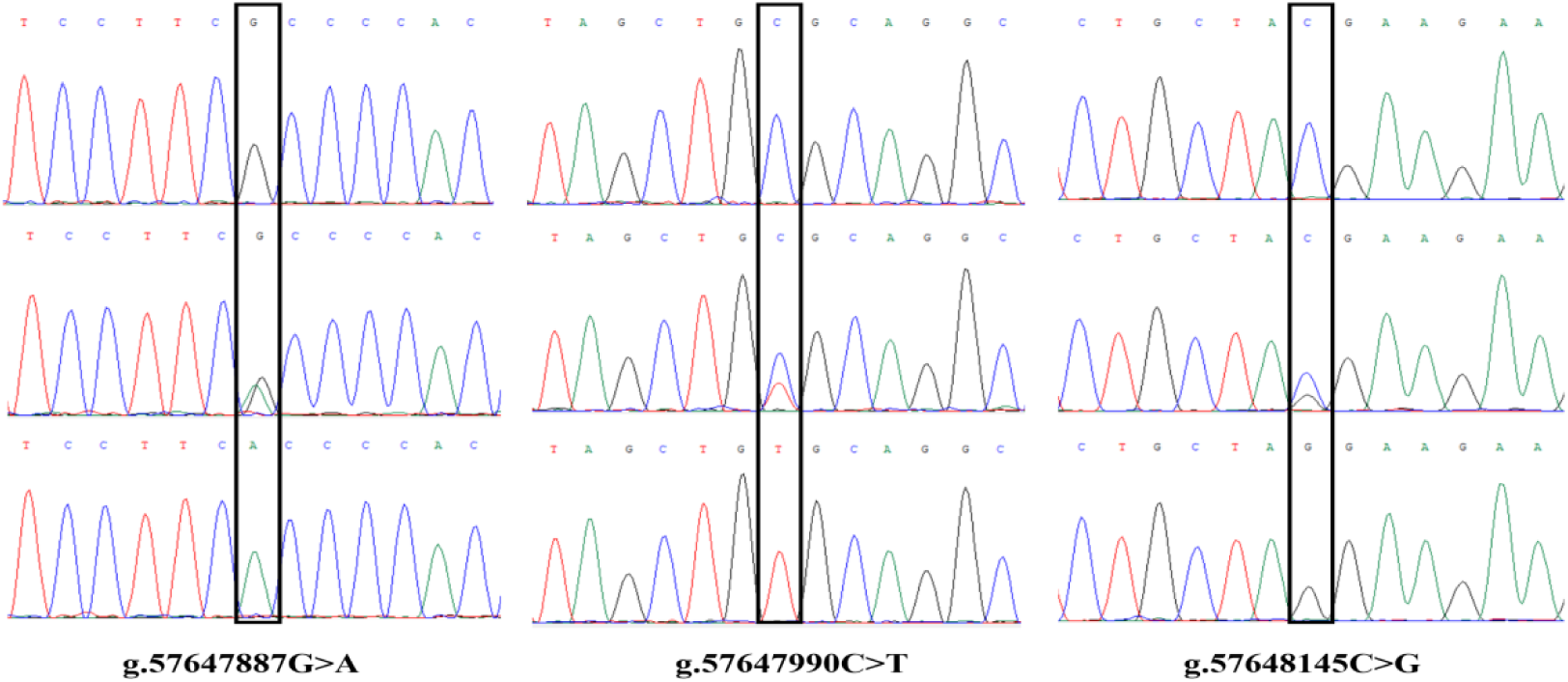
The sequence peak map of the SNPs loci in *BMP*7 gene.

### Population genetic information analysis

We counted the gene frequencies, genotype frequencies and genetic indexes of the 9 SNPs, and the results were shown in Table 2. The dominant genotypes and frequencies of the SNPs g.1884992 T>C, g.1885152 G>C and g.1887834 G>A were TT(0.68), GG(0.60), GG(0.60) respectively, and the dominant alleles and frequencies were T(0.65), G(0.73), G(0.73). The result of genetic index analysis revealed that the number of effective alleles of the SNPs g.1884992 T>C, g.1885152 G>C and g.1887834 G>A were 1.66, 1.46, and 1.46, with heterozygosity values of 0.26, 0.40, 0.40, indicating low heterozygosity. Polymorphism information content analysis revealed that the PIC values of SNPs g.1884992 T>C, g.1885152 G>C and g.1887834 G>A were 0.26, 0.32, and 0.32, representing g.1885152 G>C and g.1887834 G>A belong to intermediate polymorphism (0.25<*PIC*<0.5), and g.1884992 T>C belong to low polymorphism (*PIC<*0.25). Chi-square tests indicated that SNPs g.1884992 T>C deviates from Hardy-Weinberg equilibrium (0.05>*P*>0.01), meanwhile SNPs g.1885152 G>C and g.1887834 G>A significantly deviated from Hardy-Weinberg equilibrium (*P<*0.01).

**Table 2.** Population genetic information of *PAEP, KRT*10 and *BMP*7 SNPs in Kele pigs.

| SNPs | Genotype frequency | | | Allele frequency | | *Ne* | *PIC* | $\chi^2$ | *He* |
| --- | --- | --- | --- | --- | --- | --- | --- | --- | --- |
| g.1884992 T>C | TT (173) | TC(65) | CC (17) | T | C | 1.46 | 0.26 | 8.75 | 0.69 |
|  | 0.68 | 0.25 | 0.07 | 0.65 | 0.35 |  |  |  |  |
| g.1885152 G>C<br>/1887834 G>A | GG/GG(153) | GC/GA (65) | CC/AA(37) | G/G | C/A | 1.66 | 0.32 | 32.53 | 0.40 |
|  | 0.60 | 0.25 | 0.15 | 0.73 | 0.27 |  |  |  |  |
| g.21643703 C>T | CC(115) | CT (113) | TT(27) | T | C | 1.79 | 0.34 | 0.01 | 0.44 |
|  | 0.45 | 0.44 | 0.11 | 0.67 | 0.33 |  |  |  |  |
| g.21643714 G>A | GG(200) | GA (45) | AA(10) | G | A | 1.29 | 0.20 | 10.88 | 0.22 |
|  | 0.78 | 0.18 | 0.04 | 0.87 | 0.13 |  |  |  |  |
| g.21643741 G>A | GG(162) | GA(78) | AA(15) | G | A | 1.50 | 0.28 | 1.79 | 0.33 |
|  | 0.65 | 0.31 | 0.06 | 0.79 | 0.21 |  |  |  |  |
| g.57647887 G>A/<br>g.57647990 C>T | GG/CC(95) | GA/CT(102) | AA/TT(58) | G/C | A/T | 1.96 | 0.37 | 8.52 | 0.49 |
|  | 0.39 | 0.44 | 0.17 | 0.57 | 0.43 |  |  |  |  |
| g.57648145 C>G | CC(79) | CG(106) | GG(70) | C | G | 2.00 | 0.38 | 7.16 | 0.50 |
|  | 0.31 | 0.42 | 0.27 | 0.52 | 0.48 |  |  |  |  |

Note: *He* heterozygosity is observed, *Ne* is the effective number of the alleles, *PIC* is the polymorphic information content, $\chi^2$ -HWE is the genotype Hardy-weinberg equilibrium, $\chi^2$ -0.05=5.991, $\chi^2$ -0.01=9.210. The next tables are similar.

In the SNPs g.21643703 C>T, g.21643714 G>A and g.21643741 G>A, the dominant genotypes and frequencies were CC(0.45), GG(0.78), GG(0.65), and the dominant alleles and frequencies were C(0.67), G(0.87), G(0.79), respectively. The number of effective alleles of the SNPs were 0.44, 0.22, and 0.33, with heterozygosity values of 0.44, 0.22, 1.79 respectively, indicating low heterozygosity. The SNPs g.21643703 C>T and g.21643741 G>A exhibited intermediate polymorphism (0.25<*PIC*<0.5), and g.21643714 G>A showed low polymorphism (*PIC<*0.25). The SNPs g.21643703 C>T and g.21643741 G>A were in agreement with Hardy-Weinberg equilibrium (*P>*0.05), and g.21643714 G>A significantly deviated from Hardy-Weinberg equilibrium (*P<*0.01).

Results showed that for the SNPs g.57647887 G>A, g.57647990 C>T and g.57648145 C>G, the dominant genotypes and alleles respectively were GA(0.44), CT(0.44), CG(0.46), G(0.57), C(0.57), C(0.52). The number of effective alleles were 1.96, 1.96, and 2.00, with heterozygosity values of 0.49, 0.49, 0.50, indicating low heterozygosity. All SNPs exhibited intermediate polymorphism (0.25<*PIC*<0.5), and deviated from Hardy-Weinberg equilibrium (*P*>0.05).

### Haplotype and genotype analyses, Linkage Disequilibrium analysis of *PAEP*

As shown in Table 3, there were 3 haplotypes and 6 diplotypes in *PAEP.* The dominant haplotype was H1(TGG, 0.53). The dominant diplotype was H1H1 (TTGGGG, 0.37). For the SNPs g.1884992 T>C and g.1885152G>C (Table 4), the D’ values was 0.618, and r^2^ was 0.023. The D’ value and r^2^ between g.1884992 T>C and g.1887834 G>A were respectively 0.618, 0.023. The D’ value and r^2^ between g.1885152 G>C and g.1887834 G>A were both 1.000, indicating a complete linkage (D’ > 0.85,r^2^ > 0.33). There was no strong linkage disequilibrium effect between other SNPs.

**Table 3.**
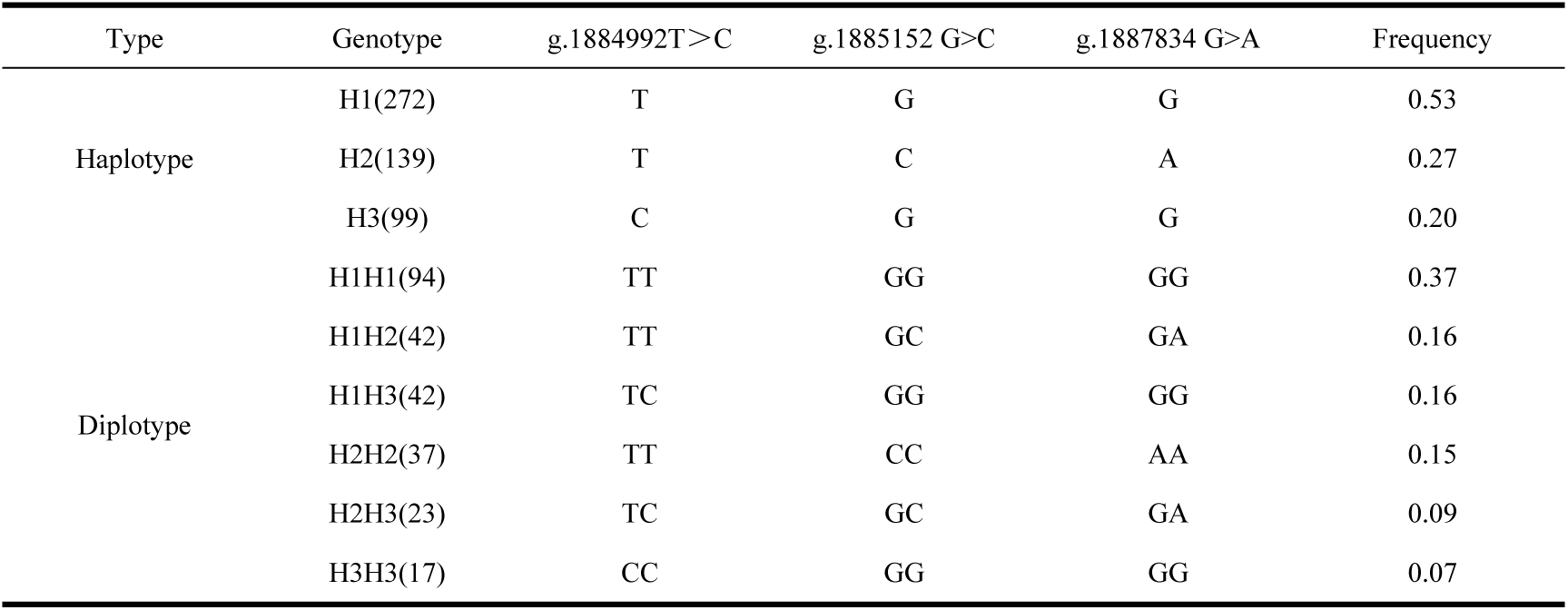
Analyses of haplotype and diplotype of *PAEP* gene in Kele pigs.

**Table 4.** Linkage disequilibrium analysis of *PAEP* gene in Kele pigs.

| SNPs | g.1884992 T>C | g.1885125G>C | g.1887834 G>A |
| --- | --- | --- | --- |
| g.1884992 T>C | - | 0.62 | 0.62 |
| g.1885152 G>C | 0.02 | - | 1.00 |
| g.1887834 G>A | 0.02 | 1.00 | - |

Note: The upper triangle are the D' values and the lower triangle are the r<sup>2</sup> values. The next tables are same.

### Haplotype and genotype analyses, Linkage Disequilibrium analysis of *KRT*10

It was observed that four haplotypes and ten diplotypes were in *KRT*10 (Table 5). The major haplotype was H1 (CGG), with a highest frequency of 0.33. The dominant diplotype was H1H2 (CTGGGG), with a frequency of 0.24. The r^2^ values among these SNPs were all lower than 0.33, therefore there was no strong linkage disequilibrium among the three SNPs (Table 6).

**Table 5.** Analyses of haplotype and diplotype in *KRT*10 gene.

| Type | Genotype | g.21643703 C>T | g.21643714 G>A | g.21643741 G>A | Frequency |
| --- | --- | --- | --- | --- | --- |
| Haplotype | H1 (170) | C | G | G | 0.33 |
|  | H2 (167) | T | G | G | 0.33 |
|  | H3 (108) | C | G | A | 0.21 |
|  | H4 (65) | C | A | G | 0.13 |
| Diplotype | H1H1 (29) | CC | GG | GG | 0.11 |
|  | H1H2 (60) | CT | GG | GG | 0.24 |
|  | H1H3 (35) | CC | GG | GA | 0.14 |
|  | H1H4 (17) | CC | GA | GG | 0.07 |
|  | H2H2 (27) | TT | GG | GG | 0.11 |
|  | H2H3 (34) | CT | GG | GA | 0.13 |
|  | H2H4 (19) | CT | GA | GG | 0.08 |
|  | H3H3 (15) | CC | GG | AA | 0.06 |
|  | H3H4 (9) | CC | GA | GA | 0.04 |
|  | H4H4 (10) | CC | AA | GG | 0.04 |

**Table 6.** Linkage disequilibrium analysis of *KRT*10 gene in Kele pigs.

| SNPs | g.1884992 T>C | g.1885125G>C | g.1887834 G>A |
| --- | --- | --- | --- |
| g.21643703 C>T | - | 0.99 | 1.00 |
| g.21643714 G>A | 0.07 | - | 0.99 |
| g.21643741 G>A | 0.13 | 0.04 | - |

### Haplotype and genotype analyses, Linkage Disequilibrium analysis of *BMP*7

Table 7 showed that four haplotypes and nine diplotypes were in *BMP*7, H1(GCC) and H1H2(GACTGC) were the most frequent of them, with the highest rates of 0.37, 0.22 respectively. The linkage disequilibrium analyses showed the SNPs g.57647887 G>A and g.57647990 C>T had a complete linkage effect, meanwhile there was no strong linkage disequilibrium effect between the remaining SNPs (Table 8).

**Table 7.**
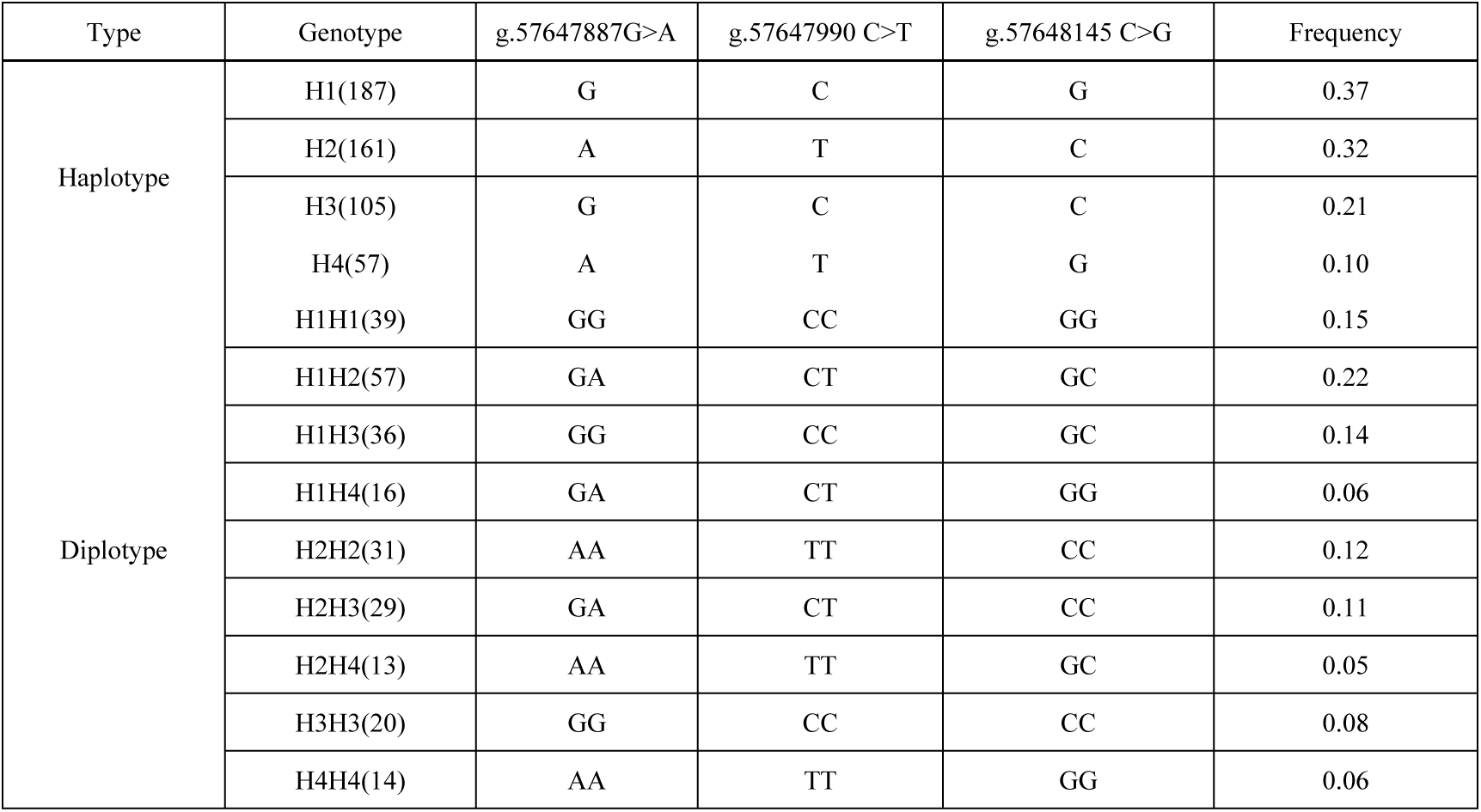
Analyses of haplotype and diplotype in *BMP*7 gene in Kele pigs.

**Table 8.** Linkage disequilibrium analysis of *BMP*7 gene in Kele pigs.

| SNPs | g.57647887 G>A | g.57647990 C>T | g.57648145 C>G |
| --- | --- | --- | --- |
| g.57647887 G>A | - | 1.00 | 0.64 |
| g.57647990 C>T | 0.97 | - | 0.64 |
| g.57648145 C>G | 0.23 | 0.22 | - |

### mRNA Secondary Structure Predictions for *PAEP* and *KRT*10 genes

We conducted mRNA secondary structure predictions for the SNPs g.1884992 T>C (4 exon) of *PAEP*, and g.21643741 G>A (5 exon) of *KRT*10, by using the online tool RNAfold (http://rna.tbi.univie.ac.at/cgi-bin/RNAWebSuite/RNAfold.cgi) (Fig. 4 and Fig.5). It is worth noticing that there were some differences in mRNA secondary structures before and after the mutations. The minimum free energy of SNPs g.1884992 T>C changed from −1276.28 kJ/mol to −1287.99 kJ/mol, indicating a decrease. In SNPs g.21643741 G>A, the minimum free energy was −2923.60 kJ/mol before mutation and −2921.69 kJ/mol after mutation, suggesting an increase.

**Fig. 4.**
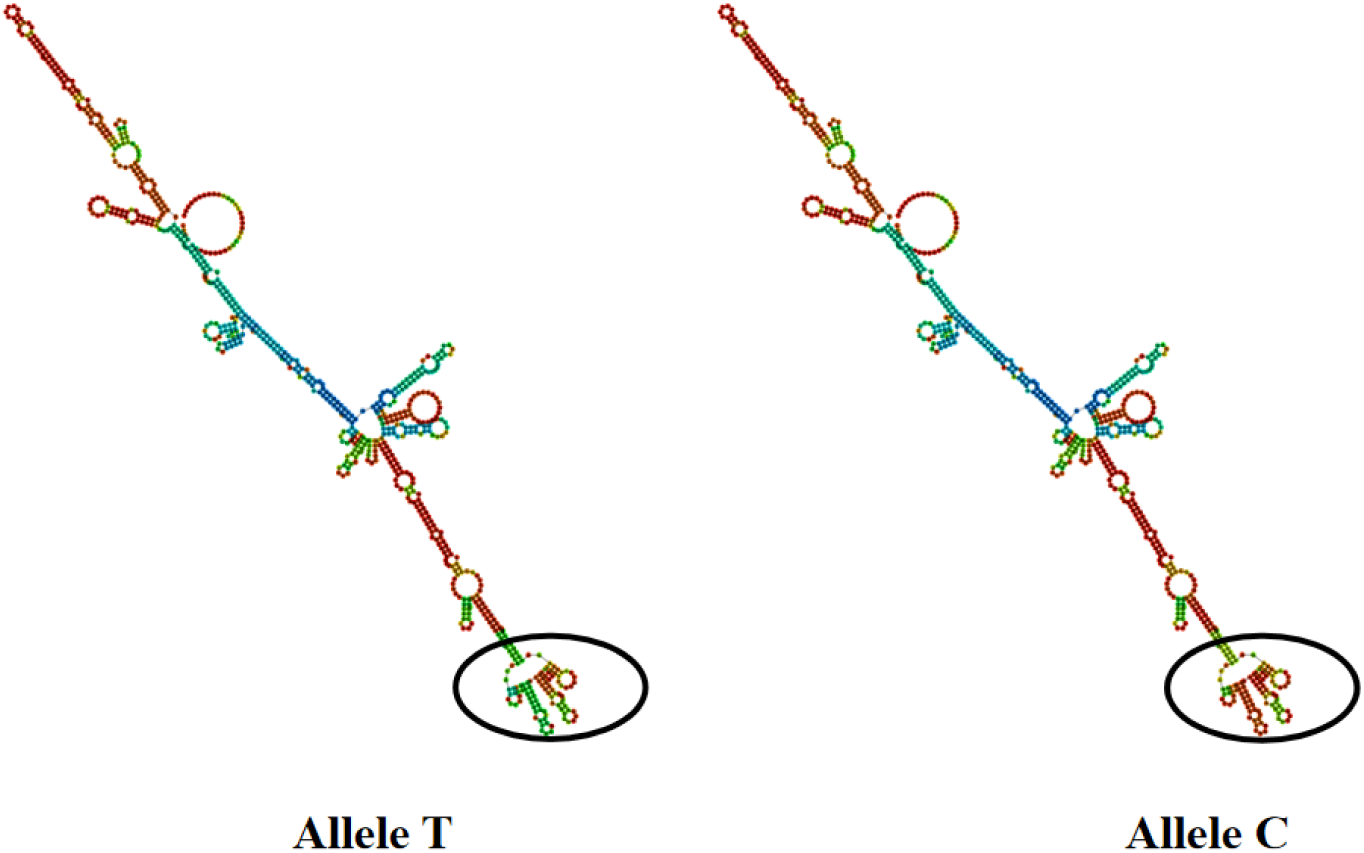
The change of mRNA secondary structure in SNPs g.1884992 T>C for *PAEP*.

**Fig. 5.**
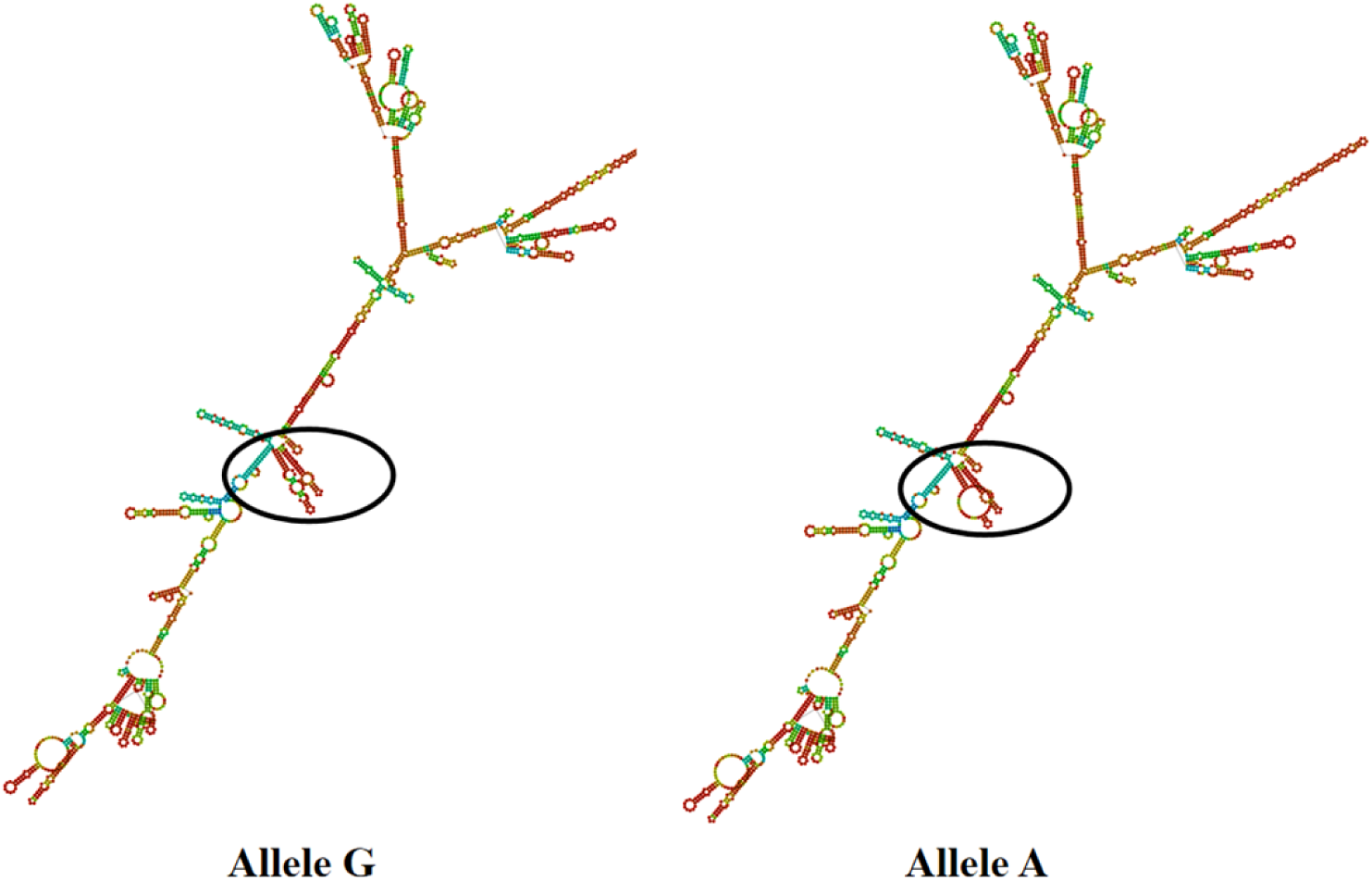
The change of mRNA secondary structure in SNPs g.21643741 G>A for *KRT*10.

### Predictions of *PAEP* Protein Secondary and Tertiary Structures

We used the online tool SOPMA (https://npsa-prabi.ibcp.fr/cgi-bin/npsa_automat.pl?page=npsa%20_sopma.html) to predict the secondary structure of the protein encoded by the *PAEP* gene in Kele pigs (Fig. 6). Before the SNPs g.1884992 T>C mutation, the percentages of α-helix, beta strand, and random coil were 12.47%, 12.98%, and 67.22%, respectively. After mutation, the percentages of α-helix, beta strand, and random coil were 12.78%, 13.12%, and 66.78%, respectively. We employed the online software SWISS-MODEL (https://swissmodel.expasy.org/interactive) and Swiss-PdbViewer software to detect the influence on entire and local tertiary structural rotein model caused by SNPs g.1884992 T>C in the Fig. 7.

**Fig. 6.**
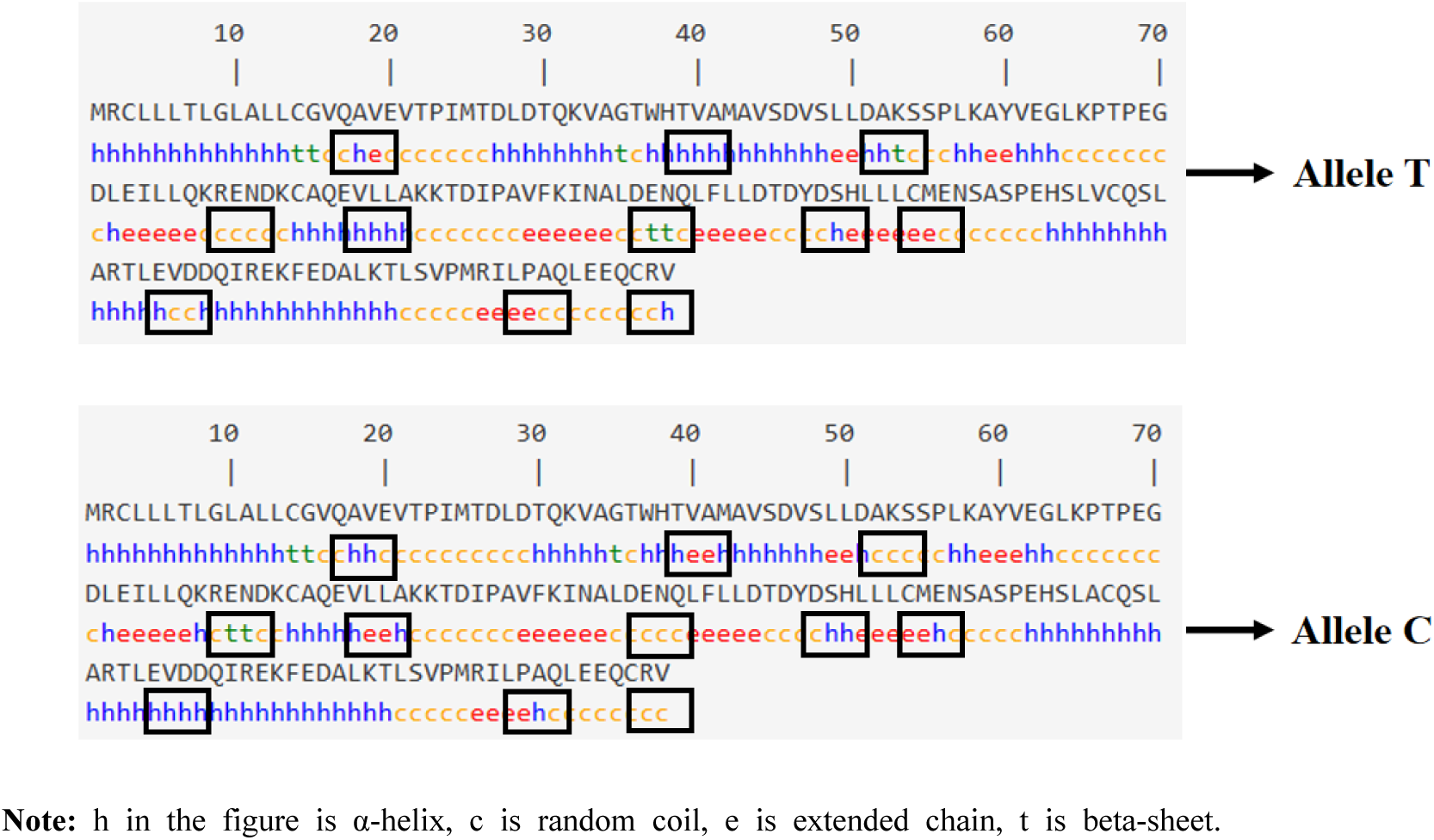
The change of secondary protein structure about SNPs g.1884992 T>C in *PAEP* gene.

**Fig. 7.**
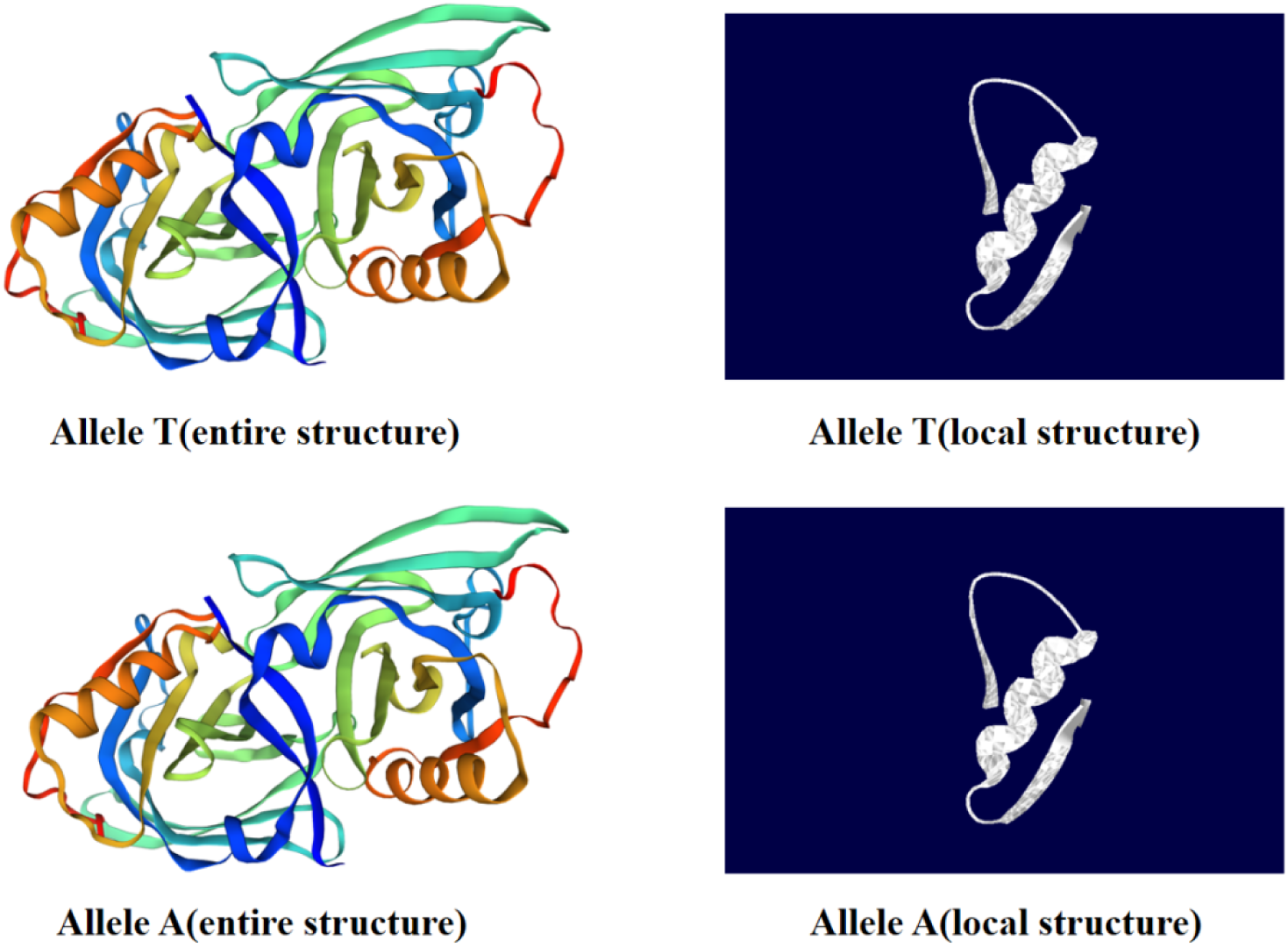
The change of entire and local tertiary protein structure about SNPs g.1884992 T>C in *PAEP* gene.

### Association between the SNPs, diplotypes of *PAEP* gene and reproductive traits in Kele pigs

The association analysis between SNPs of *PAEP* and reproductive traits was presented in Table 9. In the SNPs g.1884992 T>C, the individuals of TT genotype exhibited significantly higher total number of piglets born and number of piglets born alive compared to those of TC genotype (*P<*0.05). In the SNPs g.1885152 G>C and g.1887834 G>A, CC and AA genotypes displayed significantly higher total number of piglets born, number of piglets born alive, number of piglets weaned, and litter weight weaned than the remaining genotypes (*P<*0.05). GC genotype of SNPs g.1885152 G>C and GA genotype of SNPs g.1887834 G>A had significantly higher numbers of piglets weaned and litter weight weaned compared to GG genotypes (*P<*0.05).

**Table 9.** Association analysis between *PAEP* SNPs and reproductive traits in Kele pigs.

| SNPs | Genotype | Total number<br>of piglets born | Number of piglets<br>born alive | Litter birth<br>weight/kg | Number of piglets<br>weaned | Litter weight<br>weaned/kg |
| --- | --- | --- | --- | --- | --- | --- |
| g.1884992<br>T>C | TT(173) | 9.38±2.98<sup>a</sup> | 8.65±3.03<sup>a</sup> | 8.39±3.26 | 6.98±2.71 | 37.32±15.40 |
|  | TC (65) | 7.95±2.16<sup>b</sup> | 7.74±2.62<sup>b</sup> | 8.63±3.02 | 6.53±2.28 | 35.68±13.17 |
|  | CC(17) | 8.50±1.73<sup>ab</sup> | 7.50±1.57<sup>ab</sup> | 7.66±0.80 | 6.00±1.28 | 40.51±10.33 |
| g.1885152<br>G>C | GG(153) | 8.50±2.46<sup>b</sup> | 8.02±2.58<sup>b</sup> | 8.26±3.06 | 6.29±2.18<sup>c</sup> | 34.27±12.76<sup>c</sup> |
|  | GC(65) | 9.26±2.93<sup>b</sup> | 8.42±3.12<sup>b</sup> | 8.60±3.24 | 7.37±2.68<sup>b</sup> | 40.64±15.79<sup>b</sup> |
|  | CC(37) | 13.50±2.74<sup>a</sup> | 13.00±2.19<sup>a</sup> | 10.03±1.73 | 11.50±1.64<sup>a</sup> | 60.58±2.71<sup>a</sup> |
| g.1887834 | GG(153) | 8.50±2.46<sup>b</sup> | 8.02±2.58<sup>b</sup> | 8.26±3.06 | 6.29±2.18<sup>c</sup> | 34.27±12.76<sup>c</sup> |
| G>A | GA(65) | 9.26 ± 2.93<sup>b</sup> | 8.42 ± 3.12<sup>b</sup> | 8.60 ± 3.24 | 7.37 ± 2.68<sup>b</sup> | 40.64 ± 15.79<sup>b</sup> |
|  | AA(37) | 13.50 ± 2.74<sup>a</sup> | 13.00 ± 2.19<sup>a</sup> | 10.03 ± 1.73 | 11.50 ± 1.64<sup>a</sup> | 60.58 ± 2.71<sup>a</sup> |

Note: different shoulder letters between two pairs in the same column mean significant difference ( $P < 0.05$ ), while others mean in significant difference ( $P > 0.05$ ). The next table is similar.

The correlations between diplotypes of *PAEP* and reproductive traits in Kele pigs were shown in Table 10. For total number of piglets born, individuals with the H3H3 diplotypes revealed a significantly higher value than those of H1H1, H1H2, H1H3, H2H2, H2H3 (*P*<0.05), and H1H3 diplotypes were significantly higher than H1H1, H1H2, H2H2, H2H3 (*P*<0.05). There was no significant difference among individuals with H1H1, H1H2, H2H2, H2H3 diplotypes in total number of piglets born (*P>*0.05). In number of piglets born alive, H3H3 diplotypes were significantly higher than H1H1, H1H2, H1H3, H2H2, H2H3 (*P<*0.05), and H1H3 diplotypes were significantly higher than H2H3 (*P<*0.05). For litter birth weight, H1H2 diplotype was significantly higher than H1H1 (*P<*0.05). In the trait of number of piglets weaned, H3H3 diplotypes were significantly higher than H1H1, H1H2, H1H3, H2H2, H2H3 (*P<*0.05), and H1H3 diplotypes were significantly higher than H1H1, H1H2, H2H2, H2H3 (*P<*0.05). For litter weight weaned, H3H3 diplotypes were significantly higher than H1H1, H1H2, H1H3, H2H2, H2H3 (*P<*0.05), and H1H3 diplotypes were significantly higher than H1H1, H1H2, H2H3 (*P<*0.05).

**Table 10.**
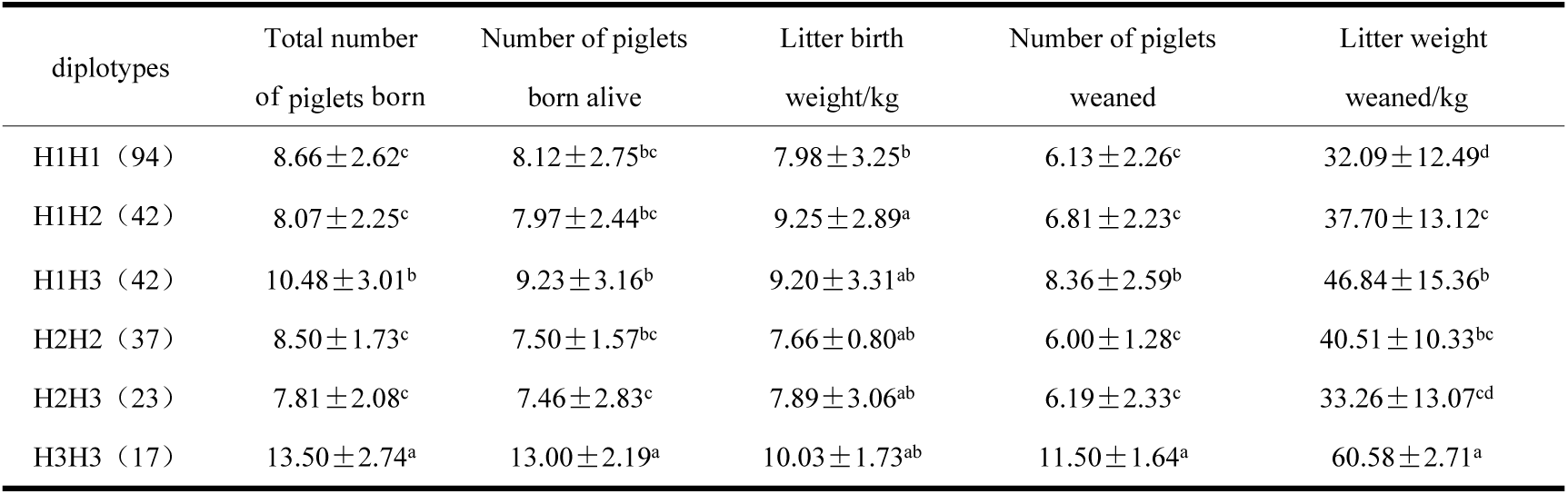
Correlation between *PAEP* diplotypes and reproductive traits in Kele pigs.

| diplotypes | Total number<br>of piglets born | Number of piglets<br>born alive | Litter birth<br>weight/kg | Number of piglets<br>weaned | Litter weight<br>weaned/kg |
| --- | --- | --- | --- | --- | --- |
| H1H1 (94) | 8.66 ± 2.62<sup>c</sup> | 8.12 ± 2.75<sup>bc</sup> | 7.98 ± 3.25<sup>b</sup> | 6.13 ± 2.26<sup>c</sup> | 32.09 ± 12.49<sup>d</sup> |
| H1H2 (42) | 8.07 ± 2.25<sup>c</sup> | 7.97 ± 2.44<sup>bc</sup> | 9.25 ± 2.89<sup>a</sup> | 6.81 ± 2.23<sup>c</sup> | 37.70 ± 13.12<sup>c</sup> |
| H1H3 (42) | 10.48 ± 3.01<sup>b</sup> | 9.23 ± 3.16<sup>b</sup> | 9.20 ± 3.31<sup>ab</sup> | 8.36 ± 2.59<sup>b</sup> | 46.84 ± 15.36<sup>b</sup> |
| H2H2 (37) | 8.50 ± 1.73<sup>c</sup> | 7.50 ± 1.57<sup>bc</sup> | 7.66 ± 0.80<sup>ab</sup> | 6.00 ± 1.28<sup>c</sup> | 40.51 ± 10.33<sup>bc</sup> |
| H2H3 (23) | 7.81 ± 2.08<sup>c</sup> | 7.46 ± 2.83<sup>c</sup> | 7.89 ± 3.06<sup>ab</sup> | 6.19 ± 2.33<sup>c</sup> | 33.26 ± 13.07<sup>cd</sup> |
| H3H3 (17) | 13.50 ± 2.74<sup>a</sup> | 13.00 ± 2.19<sup>a</sup> | 10.03 ± 1.73<sup>ab</sup> | 11.50 ± 1.64<sup>a</sup> | 60.58 ± 2.71<sup>a</sup> |

### Association between the SNPs, diplotypes of *KRT*10 gene and reproductive traits in Kele pigs

According to the result in Table 11, CC genotype of the SNPs g.21643703 C>T had significantly higher values in litter birth weight and litter weight weaned compared to those of CT and TT genotypes (*P*<0.05), which also had significantly higher number of piglets weaned compared to CT genotype (*P*<0.05). In the SNPs g.21643714 G>A, AA genotype exhibited significantly higher total number of piglets born and litter weight weaned than GA genotypes (*P*<0.05). However, there was no significant differences in number of piglets born live, litter birth weight, and number of piglets weaned in SNPs g.21643714 G>A (*P*>0.05). In the SNPs g.21643741 G>A, GG genotype showed significantly higher total number of piglets born, number of piglets born alive, number of piglets weaned, and litter weight weaned compared to AA genotypes (*P*<0.05). Meanwhile, it had significantly higher total number of piglets born compared to GA genotypes (*P*<0.05).

**Table 11.** Association analysis between *KRT*10 SNPs and reproductive traits in Kele pigs.

| SNPs | Genotypes | Total number<br>of piglets born | Number of piglets<br>born alive | Litter birth<br>weight/kg | Number of piglets<br>weaned | Litter weight<br>weaned/kg |
| --- | --- | --- | --- | --- | --- | --- |
| g.21643703<br>C>T | CC(115) | 8.83±2.74 | 8.49±2.51 | 8.88±2.69<sup>a</sup> | 7.26±2.29<sup>a</sup> | 41.33±14.22<sup>a</sup> |
|  | CT(113) | 8.75±2.57 | 8.27±2.82 | 8.07±3.42<sup>b</sup> | 6.36±2.52<sup>b</sup> | 35.29±15.35<sup>b</sup> |
|  | TT(27) | 8.56±2.81 | 7.44±3.12 | 7.03±2.9<sup>b</sup> | 6.44±3.23<sup>ab</sup> | 32.83±13.17<sup>b</sup> |
| g.21643714<br>G>A | GG(200) | 7.90±2.18<sup>ab</sup> | 7.60±1.71 | 7.81±0.84 | 6.40±1.27 | 35.57±1.95<sup>ab</sup> |
|  | GA(45) | 7.87±3.02<sup>b</sup> | 7.67±3.13 | 8.01±3.87 | 6.27±2.10 | 33.81±13.91<sup>b</sup> |
|  | AA(10) | 9.01±2.55<sup>a</sup> | 8.46±2.66 | 8.43±2.98 | 6.91±2.66 | 38.74±15.41<sup>a</sup> |
| g.21643741<br>G>A | GG(162) | 10.4±1.24<sup>a</sup> | 9.80±1.01<sup>a</sup> | 9.58±2.43 | 8.20±1.37<sup>a</sup> | 47.42±6.03<sup>a</sup> |
|  | GA(78) | 8.87±2.83<sup>b</sup> | 8.41±2.68<sup>ab</sup> | 8.39±2.53 | 7.04±2.54<sup>ab</sup> | 39.86±14.73<sup>a</sup> |
|  | AA(15) | 8.56±2.63<sup>b</sup> | 8.08±2.82<sup>b</sup> | 8.18±3.38 | 6.52±2.57<sup>b</sup> | 35.84±15.20<sup>b</sup> |

**Table 12.** Correlation between *KRT*10 diplotypes and reproductive traits in Kele pigs.

| diplotypes | Total number<br>of piglets born | Number of piglets<br>born alive | Litter birth<br>weight/kg | Number of piglets<br>weaned | Litter weight<br>weaned/kg |
| --- | --- | --- | --- | --- | --- |
| H1H1 (29) | 9.00±2.28<sup>ab</sup> | 8.90±2.57<sup>ab</sup> | 10.33±3.47<sup>a</sup> | 7.72±2.07<sup>ab</sup> | 46.58±15.2<sup>a</sup> |
| H1H2 (60) | 8.65±2.50<sup>ab</sup> | 8.10±2.63<sup>ab</sup> | 7.64±3.01<sup>bc</sup> | 6.15±2.75<sup>b</sup> | 33.87±15.79<sup>b</sup> |
| H1H3 (35) | 9.97±3.05<sup>ab</sup> | 9.37±2.56<sup>ab</sup> | 8.83±1.52<sup>ab</sup> | 7.60±2.89<sup>ab</sup> | 41.30±14.39<sup>ab</sup> |
| H1H4 (17) | 6.65±1.90<sup>d</sup> | 6.47±1.91<sup>c</sup> | 7.67±2.23<sup>bc</sup> | 5.94±1.71<sup>b</sup> | 32.23±17.33<sup>b</sup> |
| H2H2 (27) | 8.56±2.81<sup>b</sup> | 7.44±3.12<sup>bc</sup> | 7.03±2.90<sup>c</sup> | 6.44±3.23<sup>ab</sup> | 32.83±13.17<sup>b</sup> |
| H2H3 (34) | 8.41±2.16<sup>bc</sup> | 7.97±2.56<sup>ab</sup> | 8.37±2.98<sup>ab</sup> | 6.65±2.15<sup>ab</sup> | 38.90±16.08<sup>ab</sup> |
| H2H4 (19) | 9.68±3.30<sup>ab</sup> | 9.37±3.64<sup>ab</sup> | 8.92±5.03<sup>ab</sup> | 6.53±2.44<sup>ab</sup> | 33.28±11.81<sup>b</sup> |
| H3H3 (15) | 10.40±1.24<sup>a</sup> | 9.80±1.01<sup>a</sup> | 9.58±2.43<sup>ab</sup> | 8.20±1.37<sup>a</sup> | 47.42±6.03<sup>a</sup> |
| H3H4 (9) | 6.33±2.18<sup>d</sup> | 6.33±2.18<sup>c</sup> | 6.72±3.38<sup>c</sup> | 6.33±2.18<sup>b</sup> | 37.92±11.15<sup>ab</sup> |
| H4H4 (10) | 7.90±2.18<sup>cd</sup> | 7.60±1.71<sup>bc</sup> | 7.81±0.84<sup>b</sup> | 6.40±1.27<sup>ab</sup> | 35.57±1.95<sup>b</sup> |

As shown in Table 11, for total number of piglets born, the values of H3H3 diplotypes was significantly higher than those of H1H4, H2H2, H2H3, H3H4, H4H4 (*P<*0.05), and H1H1, H1H2, H1H3, H2H4 diplotypes were significantly higher than H1H4, H3H4, H4H4 (*P<*0.05). Regarding number of piglets born alive, H3H3 diplotypes was significantly higher than H1H4, H2H2, H3H4, H4H4 (*P<*0.05), and H1H1, H1H2, H1H3, H2H3, H2H4 diplotypes were significantly higher than H1H4, H3H4 (*P<*0.05). For litter birth weight, H1H1 diplotypes was significantly higher than H1H2, H1H4, H2H2, H3H4, H4H4 (*P<*0.05), and H1H3, H2H3, H2H4, H3H3, H4H4 diplotypes were significantly higher than H2H2, H3H4 (*P<*0.05). In the trait of number of piglets weaned, H3H3 diplotypes was significantly higher than H1H2, H1H4, H3H4 (*P<*0.05). For litter weight weaned, H1H1and H3H3 diplotypes were significantly higher than H1H2, H1H4, H2H2, H2H4, H4H4 (*P<*0.05).

### Association between the SNPs, diplotypes of *BMP*7 gene and reproductive traits in Kele pigs

For the *BMP*7 gene (Table 13), individuals with the AG genotype of SNPs g.57647887 G>A had significantly higher total number of piglets born and number of piglets born live compared to AA genotypes (*P*<0.05). In SNPs g.57647990 C>T, CT genotype exhibited significantly higher total number of piglets born and number of piglets born live than TT genotypes (*P*<0.05). GG genotype displayed significantly higher litter weight weaned compared to CC genotypes in SNPs g.57648145 C>G (*P*<0.05).

**Table 13.** Association analysis between *BMP*7 SNPs and reproductive traits in Kele pigs.

| SNPs | Genotypes | Total number<br>of piglets born | Number of piglets<br>born alive | Litter birth<br>weight/kg | Number of piglets<br>weaned | Litter weight<br>weaned/kg |
| --- | --- | --- | --- | --- | --- | --- |
| g.57647887<br>G>A | GG(95) | 8.00±2.60<sup>ab</sup> | 7.26±2.75<sup>ab</sup> | 7.68±2.45 | 6.26±2.02 | 36.49±13.74 |
|  | GA(102) | 9.09±2.12<sup>a</sup> | 8.26±2.22<sup>a</sup> | 8.07±3.15 | 6.86±2.76 | 36.68±13.21 |
|  | AA(58) | 8.15±3.34<sup>b</sup> | 7.77±3.45<sup>b</sup> | 7.65±3.27 | 6.15±2.71 | 34.56±16.99 |
| g.57647990<br>C>T | CC(95) | 8.00±2.60<sup>ab</sup> | 7.26±2.75<sup>ab</sup> | 7.68±2.45 | 6.26±2.02 | 36.49±13.74 |
|  | CT(102) | 9.09±2.12<sup>a</sup> | 8.26±2.22<sup>a</sup> | 8.07±3.15 | 6.86±2.76 | 36.68±13.21 |
|  | TT(58) | 8.15±3.34<sup>b</sup> | 7.77±3.45<sup>b</sup> | 7.65±3.27 | 6.15±2.71 | 34.56±16.99 |
| g.57648145<br>C>G | CC(79) | 8.71±2.87 | 8.33±2.93 | 7.72±2.76 | 6.13±2.21 | 34.31±13.05<sup>b</sup> |
|  | CG(106) | 8.5±2.22 | 7.36±2.56 | 7.81±3.00 | 6.50±2.62 | 35.25±14.31<sup>ab</sup> |
|  | GG(70) | 8.26±2.86 | 7.9±2.52 | 8.09±2.95 | 7.00±2.55 | 40.61±14.08<sup>a</sup> |

The association analysis of *PAEP* gene diplotypes with reproductive traits in Kele pigs was analyzed in Table 14. For total number of piglets born, H1H1, H1H2, H1H3, H1H4, H2H2, H2H3, H3H3, H4H4 diplotypes were significantly higher than H2H4 (*P<*0.05). In the trait of number of piglets born alive, H1H1, H1H2, H1H4, H2H2, H2H3, H3H3, H4H4 diplotypes were significantly higher than H2H4 (*P<*0.05). For litter birth weight, H4H4 diplotypes were significantly higher than H1H4, H2H4, H3H3 (*P<*0.05). In the trait of number of piglets weaned, H1H1, H1H2, H1H3, H1H4, H2H2, H2H3, H3H3, H4H4 diplotypes were significantly higher than H2H4 (*P<*0.05). For litter weight weaned, H1H1, H1H2, H1H3, H1H4, H2H2, H2H3, H3H3, H4H4 diplotypes were significantly higher than H2H4 (*P<*0.05).

**Table 14.** Correlation between *BMP*7 diplotypes and reproductive performance in Kele pigs.

| diplotypes | Total number<br>of piglets born | Number of piglets<br>born alive | Litter birth<br>weight/kg | Number of piglets<br>weaned | Litter weight<br>weaned/kg |
| --- | --- | --- | --- | --- | --- |
| H1H1(39) | 8.36±2.92<sup>a</sup> | 7.93±2.54<sup>a</sup> | 8.04±2.44<sup>ab</sup> | 7.21±1.85<sup>a</sup> | 42.49±11.89<sup>a</sup> |
| H1H2(57) | 9.22±1.89<sup>a</sup> | 8.17±2.16<sup>a</sup> | 8.13±2.92<sup>ab</sup> | 7.17±2.58<sup>a</sup> | 37.67±12.37<sup>a</sup> |
| H1H3(36) | 7.50±2.11<sup>a</sup> | 6.08±2.65<sup>ab</sup> | 7.72±2.77<sup>ab</sup> | 5.67±1.97<sup>a</sup> | 33.18±15.32<sup>a</sup> |
| H1H4(16) | 7.33±3.14<sup>a</sup> | 7.00±2.68<sup>a</sup> | 6.75±4.27<sup>b</sup> | 6.00±4.98<sup>a</sup> | 31.38±23.10<sup>a</sup> |
| H2H2(31) | 8.40±3.44<sup>a</sup> | 7.90±3.60<sup>a</sup> | 7.69±2.59<sup>ab</sup> | 6.50±2.40<sup>a</sup> | 36.22±16.11<sup>a</sup> |
| H2H3(29) | 9.33±2.17<sup>a</sup> | 8.89±2.14<sup>a</sup> | 8.37±3.40<sup>ab</sup> | 6.33±2.28<sup>a</sup> | 35.92±11.53<sup>a</sup> |
| H2H4(13) | 4.00±1.77<sup>b</sup> | 4.00±2.42<sup>b</sup> | 1.70±1.28<sup>c</sup> | 1.00±1.56<sup>b</sup> | 4.50±8.00<sup>b</sup> |
| H3H3(20) | 8.20±2.78<sup>a</sup> | 8.20±2.78<sup>a</sup> | 6.61±1.21<sup>b</sup> | 5.00±1.33<sup>a</sup> | 27.60±5.19<sup>a</sup> |
| H4H4(14) | 9.00±2.31<sup>a</sup> | 9.00±2.31<sup>a</sup> | 10.43±3.55<sup>a</sup> | 7.00±2.31<sup>a</sup> | 41.33±8.86<sup>a</sup> |

## Discussion

*PAEP* is a protein initially described in reproduction. It not only influence the capacitation, acrosome reaction, sperm-oocyte binding and immune system suppression during the establishment of pregnancy [32], but also is known to be expressed in female-specific tumours, such as breast [33], endometrial [34], ovarian and cervical cancer [35]. And research has shown that *PAEP* has three subtypes, which play important roles in maintaining, initiating, and prolonging the uterine environment for pregnancy and can be tested for use in assisted reproductive technology projects [36] [37]. In this study, we detected the SNPs g.1884992 T>C (exon 4), g.1885152 G>C (intron 4) and g.1887834 G>A (intron 4) in *PAEP*. The low heterozygosity suggests limited genetic variation and low diversity in this population [38]. This phenomenon may be attributed to the fact that the experimental materials originate from a specific locality, with minimal hybridization with other breeds from different regions, resulting in relatively stable genetic materials. Three SNPs all exhibited intermediate polymorphism (0.25<*PIC*<0.5), thereby offering valuable genetic information that can maintain the adaptability and survival capability of the population. The SNPs g.1884992 T>C deviated from Hardy-Weinberg equilibrium (0.01<*P*<0.05), while g.1885152 G>C and g.1887834 G>A significantly deviated from Hardy-Weinberg equilibrium (*P*<0.01), suggesting that gene distribution may be influenced by natural selection, drift, migration, and other factors [39]. The SNP of g.1884992 T>C was in the fourth extron located in the gene’s coding sequence region, which may alter the protein’s conformation. The prediction result of mRNA secondary structure showed that the free energy of mRNA secondary structure was reduced and the structural stability was enhanced after the mutation. The mRNA synthesizes proteins using itself as a template, and changes in mRNA may have an impact on the structure and function of proteins. We made further predictions of the secondary and tertiary structures of the PAEP protein to test that. The result showed that the secondary structure of the protein encoded by the SNPs g1884992T>C changed, and the tertiary structure was similar. The side chains of valine (V) and alanine (A) are both hydrophobic and have similar sizes, which may be the reason why their protein tertiary structures are similar. [40] [41]. However, there are still significant differences in their biological functions and chemical properties, and their functional expression may change in the reproductive traits of Kele pigs. The significant differences in total litter size, live litter size, weaned piglets, and weaned litter weight after the mutation may be the result of changes in protein structure and function. The SNPs g. 1885152G>C and g. 1887834G>A are both located in the fourth intron.

Although mutations at this site do not cause changes in the coding amino acids, they may play a role by regulating different gene expression levels and potentially affecting gene transcription activity, thereby altering gene function [42]. The results of the correlation analysis showed that the two SNPs had an impact on reproductive traits such as total litter size, live litter size, and weaned piglet size in Kele pigs. In this study, the individuals with CC genotype of SNPs g. 1885152 G>C and AA genotype of SNPs g.1887834 G>A had significantly higher total number of piglets born, number of piglets born alive, number of piglets weaned, and litter weight weaned than other genotypes, indicating a favorable genotype. The analysis of diplotype H3H3 (CCGGGG) individuals showed that total number of piglets born, number of piglets born alive, number of piglets weaned, and litter weight weaned were significantly higher than H1H1 (TTGGG), H1H2 (TTGCGA), H1H3 (CTGGG), H2H2 (TTCCAA), and H2H3 (TCGCGA) (P<0.05), indicating a favorable diplotype. The diplotype H3H3 (CCGGGG) can serve as a reference for molecular marker assisted selection in total number of piglets born, number of piglets born alive, number of piglets weaned, and litter weight weaned.

As genes encoding important proteins in keratinocytes, *KRT*10 and its homologues have not been reported to exhibit polymorphism in reproduction traits of livestock and poultry. But existing research has shown they are closely associated with the reproductive organs. Smedts et al. [43] detected the expression of K1, K6, K15, K16, and K20 in normal cervical epithelium, squamous metaplasia, different grades of cervical intraepithelial neoplasia, as well as single specific antibody squamous cell carcinoma and cervical adenocarcinoma. The study by RicciardelliI et al. [44] showed that K5 can be used to predict the prognosis of serous ovarian cancer and identify cancer cells resistant to chemotherapy. Wang et al. [45] found that K17 may be an important molecular marker for predicting EOC carcinogenesis, progression, and prognosis. There were three SNPs in *KRT*10, including g.21643703 C>T (intron 4), g.21643714 G>A (intron 4) and g.21643741 G>A (exon 5). The SNPs g.21643714 G>A showing low polymorphism (*PIC*<0.25). It indicated the variant alleles or genotypes of the *KRT*10 gene in the Koala pig population are relatively stable, with fewer variations, potentially resulting in lower adaptability to environmental changes. Chi-square tests showed that the genotyping in SNPs g.21643703 C>T and g.21643741 G>A were in Hardy-Weinberg equilibrium (*P*>0.05), which meant that for the two SNPs in the population, the frequencies of genotypes and alleles remained constant across generations, indicating the absence of evolutionary influences such as natural selection, mutation, migration, and genetic drift. We analyzed the molecular biological structure encoded by the SNPs g. 21643741 G>A in the fifth exon. Before and after the mutation, the amino acids enconded by SNPs g.21643741 G>A both were lysine (K), indicating a synonymous mutation. Synonymic mutation refers to the phenomenon of base replacement without causing changes in amino acid types. Kimura [46] previously believed that synonymous mutations have no impact on biological adaptability and are “neutral” during evolution. However, the research of Chamary et al. [47], showed that there is codon bias associated with synonymous codons in organisms, and synonymous mutations can cause certain functional changes in the organism. Synonymic mutations affect gene expression by altering mRNA stability, secondary structure, splicing process, and translation dynamics [48]. In addition, synonymous mutations also have an impact on protein translation rate, stability, correct folding, and other processes [49]. The prediction result of the mRNA secondary structure found that the structure changed and the structural stability weakened after mutation. In this study, the significant differences in total number of piglets born, number of piglets born alive, number of piglets weaned, and litter weight weaned of the SNPs g. 21643741 G>A may be the result of synonymous mutations. Individuals with the GG genotype in SNPs 21643741 G>A had significantly higher total number of piglets born, number of piglets born alive, number of piglets weaned, and litter weight weaned than those with the AA genotype (*P*<0.05), and significantly higher total number of piglets born than those with the GA genotype (*P*<0.05). It was preliminarily identified as a favorable genotype. Molecular breeding techniques can be latterly used to enhance the selection intensity of this locus and achieve homozygous genotype GG as soon as possible to improve reproductive traits. The analysis of the diplotypes showed that individuals with H1H1 (CCGGGGG) had higher litter birth weight, and was the optimal diplotype for increasing litter birth weight. The individuals of H3H3 (CCGGAA) diplotype reached significant levels in all traits, indicating a favorable diplotype and serving as a reference for molecular marker assisted selection.

Existing research indicates that the *BMP*7 gene has significant implications for improving reproductive traits in livestock and poultry. Xu et al. [50] found that the mRNA expression level of *BMP*7 in the follicles of multi lamb ewes of Hu sheep was significantly higher than that of single lamb individuals. Zhang et al. [51] found that changes involving transcription factors such as <u>USF1</u>, <u>USF2</u> and INMS1 in the *BMP*7 <u>promoter region</u> might be involved in greater sheep prolificacy. In the study by Yin et al. [52], the SNPs c.1569 A>G in *BMP*7 had a significant impact on some reproductive traits of sows in large white pig populations, which had some similarities with the results of this study. We found the SNPs g.57647887 G>A, g.57647990 C>T and g.57648145 C>G in *BMP*7, all in intron 3. All SNPs exhibited low heterozygosity, intermediate polymorphism (*PIC*<0.25), and deviated from Hardy-Weinberg equilibrium (0.01<*P*<0.05). The SNPs g.57647887 G>A and g.57647990 C>T exhibited significant levels on total number of piglets born and number of piglets born alive (*P*<0.05), and the individuals with GA genotype and CT genotype had higher values than others. The SNPs g.57648145 C>G significantly influenced litter weight weaned (*P*<0.05), and the individuals with GG genotype had higher values than others. Individuals with the H4H4 diplotypes exhibited significant effects on total number of piglets born, number of piglets born alive, litter birth weight, number of piglets weaned, and litter weight weaned, indicating it can be a favorable diplotypes.

## Conclusion

In this study, we totally indentified nine SNPs in *PAEP*, *KRT*10, and *BMP*7 genes. Through a comprehensive analysis, it was observed that the H3H3 (CCGGGG) diplotype of *PAEP* and the H3H3 (CCGGAA) diplotype of *KRT*10 could be used as a potential genetic marker to improve total number of piglets born, number of piglets born alive, number of piglets weaned, and litter weight weaned. Additionally, the H4H4 (AATTGG) diplotype of *BMP*7 could be used as a molecular marker for assisting in the selection of total number of piglets born, number of piglets born alive, litter birth weight, number of piglets weaned, and litter weight weaned. Consequently, the *PAEP*, *KRT*10, and *BMP*7 were all promisingly as candidate genes for the reproductive traits of Kele pigs.

## Data availability

The data that support this study will be shared upon reasonable request to the corresponding author.

## Conflict of interest

No conflict of interest exists in the submission of this article, and the article is approved by all authors for publication. I would like to declare on behalf of my co-authors that the work described is original research that has not been published previously and is not under consideration for publication elsewhere, in whole or in part.

## Declaration of funding

This research was jointly funded by the Guizhou Provincial Science and Technology Plan Project (QKHPTRC[2021]5630), Guizhou Provincial Swine Industry Technology System Construction (GZSZCYJSTX-03), and National Key Research and Development Program (2022YFD1100308-01).

## Acknowledgements

The authors gratefully acknowledge the Hezhang County Farm for providing the experimental animals and Shenggong Biotechnology Co., Ltd for gene sequencing.

## Author contribution

Yong Zhao: experimental operation, data curation, validation, formal analysis, roles/Writing-original draft. Chunyuan Wang: experimental operation, data curation, validation, investigation. Yan Wu: experimental operation, data curation, roles/Writing-original draft. Jin Xiang: data curation, formal analysis, project administration. Yiyu Zhang: conceptualization, methodology, funding acquisition.

## Notes

### Competing Interest Statement

The authors have declared no competing interest.

